# A Re-evaluation of Phylogenomic Data Reveals that Current Understanding in Wheat Blast Population Biology and Epidemiology is Obfuscated by Oversights in Population Sampling

**DOI:** 10.1101/2022.12.19.521104

**Authors:** Mark L. Farman, Joao P. Ascari, Mostafa Rahnama, Emerson M. Del Ponte, Kerry F. Pedley, Sebastián Martinez, José Maurício C. Fernandes, Barbara Valent

## Abstract

Wheat blast, caused by the *Triticum* lineage of *Pyricularia oryzae* (PoT), is a serious disease that first emerged in Brazil and quickly spread to neighboring countries. The recent appearance of this disease in Bangladesh and Zambia highlights a need to understand the population biology and epidemiology of the disease so as to mitigate pandemic outbreaks. Current knowledge in these areas is largely based on analyses of wheat blast isolates collected in Brazil, and their comparison with isolates from non-wheat, endemic grasses. Those studies concluded that wheat blast is caused by a highly diverse *P. oryzae* population that lacks host specificity and, as a result, undergoes extensive gene flow with populations infecting non-wheat hosts. Additionally, based on genetic similarity between wheat blast and isolates infecting *Urochloa* species, it was proposed that the disease originally emerged via a host jump from this grass, and the widespread use of *Urochloa* as a pasture grass likely plays a central role in wheat blast epidemiology. Inconsistencies with earlier phylogenetic studies prompted us to re-analyze the Brazilian data in the context of a comprehensive, global, phylogenomic dataset. We now show that the seminal studies failed to sample the *P. oryzae* populations normally found on endemic grasses and, instead, repeatedly sampled PoT and *P. oryzae Lolium* (PoL) members that happened to be present in these hosts. The resulting lack of accurate and representative information about the grass-infecting populations in Brazil means that current conclusions about wheat blast’s evolution, population biology and epidemiology are unsubstantiated and could be equivocal.

## INTRODUCTION

Wheat blast (also known as brusone) is a serious disease of wheat caused by the fungus *Pyricularia oryzae*. Having recently spread to Asia and Africa (Malaker et al. 2016; Tembo et al. 2020), it now poses a serious threat to global wheat production (Ceresini et al. 2018; Cruz and Valent 2017; Singh et al. 2021). As a recently emerged disease that first surfaced in Paraná state, Brazil in 1985 (Igarashi et al. 1986), wheat blast promises to be a valuable model for understanding how new diseases evolve; how newly-evolved populations are structured and change over time; how they interact with endemic *P. oryzae* populations in the field; and, then, finally, what are the population genetic consequences of invading new continents. Intimate knowledge of pathogen population structure/genetics also has practical benefits in disease control as it can provide deep insights into disease epidemiology, and underpins a solid theoretical foundation for the development of diagnostic tools and quarantine guidelines. Here, it should go without saying that accurate inference of population structure is absolutely critical to success in these endeavors (Milgroom 2017).

Despite a long history of research in blast diseases (most notably rice blast), the overall low level of sequence divergence between *P. oryzae* isolates (~1 %) prevented significant progress in understanding its population structure until the development of hypervariable molecular markers (simple sequence repeats - SSRs), and next generation sequencing. Ceresini and coworkers were pioneers with these technologies and used SSRs (Maciel et al. 2014), multilocus sequencing (Castroagudín et al. 2016), and eventually whole genome sequencing (Ceresini et al. 2018, 2019) to characterize the wheat blast population. They showed that fungal isolates causing wheat blast are phylogenetically distinct from those found on rice, *Digitaria, Eleusine, Setaria* and other grasses, and renamed the population as a new species, *Pyricularia graminis tritici* (Pygt) to reflect this fact (Castroagudín et al. 2016; Ceresini et al. 2019). More recently, however, evidence of gene flow (Gladieux et al. 2018) among other concerns (Valent et al. 2019), has raised this conclusion into doubt.

Early population studies of wheat blast using simple sequence repeats (SSRs) found these markers to be in equilibrium, while at the same time clonal lineages were identified. This implied that the population has a mixed reproductive mode, where sexual cycles generate diversity and well-adapted clones are then propagated vegetatively (Maciel et al. 2014). Ceresini and coworkers then compared wheat blast strains with fungal isolates found on neighboring grasses and weeds. Using SSRs, and then genome sequence data, they found that many grass-infecting isolates exhibited a high degree of genetic similarity to isolates found on wheat (Castroagudin et al. 2017; Ceresini et al. 2018, 2019). Also detected was evidence for significant gene flow between the grass- and wheat-infecting populations, with the predominant migration being from the former to the latter (Castroagudin et al. 2017). When combined with the observations that several isolates from grasses were capable of causing disease on wheat in inoculation assays; virulence phenotypes were often shared between the wheat and non-wheat host groups (Castroagudin et al. 2017); and the discovery of inter-fertility between isolates from wheat and other Poaceae (Bruno and Urashima 2001; Galbieri and Urashima 2008; Urashima et al. 1993); this led Ceresini and coworkers to propose that the wheat blast fungus mates preferentially on non-wheat hosts, producing an ascospore population with high diversity which then infects nearby wheat (Ceresini et al., 2018, 2019). An extended conclusion from these findings was that wheat blast (Pygt) is not a wheat-specialized pathogen so that *P. oryzae* - being predominantly host-specialized - is not a good model for studying Pygt biology (Ceresini et al., 2018, 2019).

With regard to the grass-infecting populations, significant focus has been placed on the one infecting *Urochloa* because the wheat blast pathogen reportedly bore the strongest similarity to isolates from this host. This led to the conclusion that a host jump by a *Urochloa* pathogen was a key step in the evolution of wheat blast (Stukenbrock and McDonald 2008). Furthermore, because *Urochloa* is widely used as a forage grass in Brazil, and is often grown in close proximity to wheat, it has been proposed that signalgrass pastures are significant inoculum reservoirs for pathogen over-wintering, and serve as a bridge that facilitates pathogen spread and gene flow between regions (Ceresini et al. 2019).

Unfortunately, if one peruses the phylogenomic data that underpin the foregoing conclusions, and considers them in the light of data produced by the broader research community, major inconsistencies quickly appear with potentially far-reaching consequences for foundational knowledge on wheat blast. Ceresini and coworkers’ earliest studies found that wheat blast isolates could be grouped in two main clades, with only one (Pygt) being clearly resolved from the rice blast pathogens (Castroagudin et al. 2016). Later, as genomic sequence data provided greater resolution, additional wheat blast isolates were brought under the Pygt umbrella, along with some of the isolates from grasses (Castroagudin et al. 2017; Ceresini et al. 2018, 2019). However, the *Pygt/P. oryzae* boundary shifted every time new data were added (compare Castroagudin et al., 2017 vs. Ceresini et al., 2018 vs. Ceresini et al., 2019). Not only is this a reason to doubt Pygt’s status as a new species but it also raises serious concerns about the reported relationship between Pygt and other grass pathogens.

When studied in a broader phylogenomic context, a majority of wheat blast isolates form a single, cohesive clade known as the *P. oryzae Triticum* lineage, or PoT (formerly *Magnaporthe oryzae Triticum*, MoT). A relatively small proportion of isolates group in another clade with isolates from *Lolium* species (PoL1 - so named because it is the first of three distinct lineages containing isolates from *Lolium*) (Gladieux et al. 2018). Recent studies, based on whole genome analysis of a comprehensive collection of isolates, show that within these lineages, isolates can be further grouped into discrete haplotypes based on distinct chromosomal configurations that arose when wheat blast/gray leaf spot first evolved via a process that involved recombination of divergent *P. oryzae* genomes in a multi-hybrid swarm (Rahnama et al. submitted). To date, 44 discrete haplotypes have been identified within PoT and 13 in PoL1 (Rahnama et al, submitted; Ascari et al. in preparation).

Critically, when analyzed together with a comprehensive collection of isolates from other Poaceae, the PoT and PoL lineages show complete phylogenetic resolution with extensive sequence divergence even from the nearest neighboring lineages (Gladieux et al. 2018). This is not true for the grass-infecting isolates collected in Brazil which, with just one exception, all grouped phylogenetically with Pygt - having even been included in the species in one iteration of the study (Ceresini et al. 2018). That virtually all grass-infecting isolates collected elsewhere around the globe group according to host-of-origin and are genetically very distinct from isolates found on wheat (Gladieux et al. 2018) led us to suspect that Ceresini and coworkers - having only collected isolates from grasses growing near diseased wheat - might have failed to sample the true endemic grass-infecting populations in Brazil, and instead, repeatedly recovered PoT (or PoL1). To test this possibility, we performed a new phylogenomic analysis that integrated their data with those generated by the broader *P. oryzae* research community - with the prediction that most of their isolates would group with PoT/PoL1. Then, to rule out the possibility that the *P. oryzae* populations found infecting grasses in S. America are just different to those found everywhere else around the globe, we used genome sequencing to survey a modest number of isolates collected from endemic grasses in Uruguay and at locations distant from wheat-growing regions in Minas Gerais, Brazil.

## MATERIAL AND METHODS

### Fungal isolates

Diseased leaves collected in our survey were placed in a dew chamber overnight to induce sporulation. Spores were then collected by brushing the conidiophores lightly with a sterile, sealed Pasteur pipette and then spread across 3% water agar. A single, germinated spore was then transferred to oatmeal agar supplemented with ampicillin (100 μg/ml). The colony was allowed to colonize whatman 3M paper squares placed on the agar surface, which were then collected, placed in glassine envelopes and dried in a containment hood for 2 d. The cultures were stored desiccated at −20°. The isolates that are the central focus of this paper are listed in Table 1.

### DNA extraction

Single spored isolates were cultured for 7 d with shaking on 10 ml of liquid complete medium (Valent et al. 1984). The mycelial ball was grabbed with forceps, patted dry on paper towels, placed in a 15 ml plastic, conical tube, frozen at −20 °C, and then freeze-dried overnight (https://youtu.be/h9TZANDMnd8). A glass rod was then used to grind the freeze-dried pellet to a fine powder against the side of the tube (https://youtu.be/RR0qwc3liEI). One milliliter of lysis buffer (10 mM Tris-HCL, pH 8.0; 10% SDS; 100 mM NaCl; 10 mM EDTA) was then added and mixed in by shaking and tube inversion. The buffer was incubated at room temperature for 15 min, after which 1.5 ml of phenol:chloroform:isoamyl alcohol was added and mixed in by shaking and tube inversion. The tube was allowed to sit at room temperature for 15 min with periodic re-mixing and then the mixture was fully emulsified by shaking before adding equal amounts to two tubes of Phase Lock gel (5PRIME, Gaithersburg, MD). The tubes were centrifuged for 20 min at XX g in a benchtop centrifuge and the supernatant was then decanted into two separate microfuge tubes. The DNA was precipitated by mixing in 0.54 volumes of room temperature isopropanol, followed by immediate centrifugation at XX g for 5 min. The pellets were rinsed with 70% ethanol and then allowed to air dry. Pellets were re-dissolved in TE + 10 μg/ml RNAse (Qiagen Corp.).

### Library Construction and Genome Sequencing

Sequence-ready libraries were prepared using the Nextera (Illumina Corp., San Diego, CA) and HyperPlus DNA kits (Roche Diagnostics, Indianapolis, IN). Nextera libraries were generated according to the manufacturer’s protocol with the lone modification being an extension of the tagmentation reaction time to 60 min. HyperPlus libraries were generated precisely according to the manufacturer’s protocol. Libraries were submitted to NovoGene for sequencing acquisition (150 bp paired-end reads) on the HiSeq2500 platform.

### Genome assembly

Raw sequence reads were quality filtered and adapters were trimmed using Trimmomatic (Bolger et al. 2014) with the following options: ILLUMINACLIP:adapters.fa:2:30:10 SLIDINGWINDOW:20:20 MINLEN:90. Filtered reads were then assembled using velvet 1.2.10 with the velvetoptimiser wrapper being used to iterate through kmer values of 89 to 129, with a step size of 2, to find the optimal assembly.

### Genome Masking and SNP calling

SNPs were called from masked genome alignments using two basic strategies: the first involved alignments to a common reference assembly, while the second utilized genomes aligned in all possible pairwise combinations, and was included to minimize information loss due to extensive presence/absence polymorphism. The basic algorithmic approaches, as implemented in the iSNPcaller package (https://github.com/drdna/iSNPcaller), involved: 1) masking repeats in all genomes; 2) performing pairwise alignments using BLASTn (-evalue 1e-20 -max_target_seqs 20000); 3) identifying SNPs that occur in uniquely aligned segments of both the reference and the query genome; 4) determining the total number of uniquely aligned nucleotide positions.

### Phylogenetic analyses

Neighbor joining trees were built by importing pairwise distance data from iSNPcaller into MEGA X (Kumar et al. 2018) and using the default parameters. Maximum likelihood analysis was performed using a SNP dataset that was filtered so that only those nucleotide positions called in every isolate were retained (i.e. no missing data). Tree building was performed using RAXML-NG-MPI (ref.) with the GTR + Gamma substitution model to generate 10 starting random trees, 10 starting parsimony trees, and then 100 bootstrap replications. The best tree was plotted using the R package *ggtree* (Yu et al. 2017). A geographic map was produced to indicate where the non-wheat isolates were collected, together with information on their hosts of origin and distance to wheat fields. The map was produced using the R packages *rnaturalearth*, *rnaturalearthhires* and *scatterpie* (Yu 2021; South 2021, 2022).

### Data and Code Availability

DNA sequences have been deposited at NCBI under BioProject PRJNA320483. Custom code used for data analyses is available at https://github.com/drdna/WheatBlast.

## RESULTS

### Phylogenetic analysis of the Ceresini isolates in the context of the global *P. oryzae* population

Ceresini and coworkers previously based on comparisons between a large number of fungal isolates from wheat and 15 isolates from seven different non-wheat hosts (*Avena, Cenchrus, Digitaria, Echinochloa, Eleusine, Elionurus, Melinis*, and *Urochloa*). Critically, all of the isolates from non-wheat hosts were collected from just two locations, both of which were immediately adjacent to diseased wheat (Figure S1, Table S1). To understand the relationship between the Brazilian isolates from non-wheat hosts and those collected elsewhere around the globe, the raw reads were downloaded and assembled into genome sequences. SNP calling was then performed using the iSNPcaller pipeline (https://github.com/drdna/iSNPcaller) and the resulting pairwise distance data were used to build a neighbor-joining tree. This revealed that all but one of the Ceresini isolates from non-wheat hosts grouped firmly with the PoT or PoL1 lineages, regardless of the host-of-origin (Figure 1). The isolates from *Cenchrus, Eleusine, Elionurus* and *Melinis* were members of PoT, as were three of the four isolates from *Urochloa*. All three isolates from *Avena* belonged to PoL1, which also included individual isolates from *Echinochloa, Digitaria* and *Urochloa*. These placements were made with a high degree of confidence because the branch lengths revealed significant divergence relative to the next closest lineages. The one exceptional isolate, Ds363, had previously been assigned as a separate species, *P. grisea*, and yet it clearly grouped with the *Echinochloa* lineage (PoEc), which was firmly positioned within the *P. oryzae* tree (Figure 1; Figure S2). Further support for these groupings came from a maximum likelihood analysis of a whole genome dataset based on 364,573 SNPs. This produced a very similar tree topology with strong bootstrap support for the clades representing PoT and PoL1.

**Figure 1.**
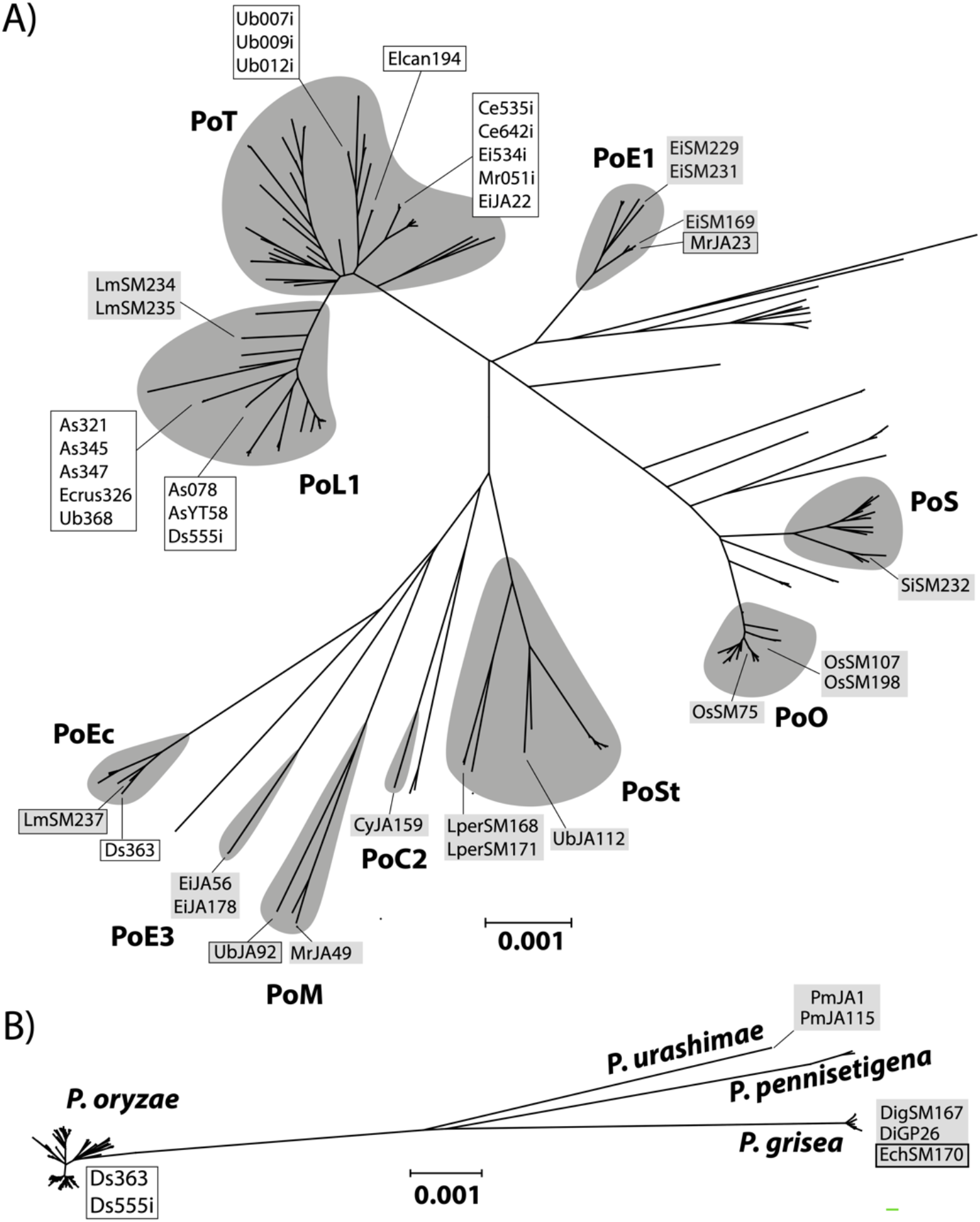
Distance trees showing the phylogenetic grouping of grass-infecting isolates from Brazil and Uruguay. A) Tree includes only *P. oryzae* isolates; B) Tree includes the four species *P. oryzae, P. grisea, P. pennisetigena* and *P. urashimae*. Ceresini isolate names are on a white background, isolates collected in the present study are shown on a gray background. Black borders are used to highlight isolates that do <u>not</u> group according to host-of-origin. The names of the relevant phylogenetic groups are shown in bold (PoC2 - Cynodon2; PoE1 - Eleusine1; PoE3 - Eleusine3; Ec - Echinochloa; PoL1 - Lolium1; PoM - Melinis; PoO - Oryza; PoS - Setaria; PoSt - Stenotaphrum. Scales show phylogenetic distance measured in substitutions per site.

Interestingly, many isolates from non-wheat hosts grouped together in the phylogeny and showed so little genome-wide divergence that they can essentially be considered clones of one another. All three of the PoT members found on *Urochloa* belonged to a single clade, and the isolates from *Cenchrus* (n=2), *Eleusine* (n=2) and *Melinis* (n=1) belonged to another. Similarly, three of four isolates from *Avena* grouped in a single clade within PoL1, along with isolates from *Echinochloa* and *Urochloa*, while the remaining *Avena* pathogen grouped with an isolate from *Digitaria*. Interestingly, although the latter clades were positioned firmly within the PoL1 lineage, neither contained isolates from *Lolium*.

### Analysis of the S. American *P. oryzae* from non-wheat hosts in Uruguay and Minas Gerais State

Because all but one of the Brazilian *P. oryzae* grouped phylogenetically with PoT/PoL1, while all isolates from *Digitaria, Eleusine*, and *Urochloa* collected elsewhere grouped according to host-of-origin, this suggested that Ceresini and coworkers had failed to sample the true, endemic grass-infecting populations in that country. However, we also considered the possibility that South American blast populations are very different to those found in the rest of the world. To distinguish between these scenarios, we examined isolates collected in S. America but from locations at varying distances from wheat fields. First, we used whole genome sequencing to analyze isolates from a number of non-wheat hosts in Uruguay where there have been no reports of wheat blast, despite being located just 750 km from Rio Grande do Sul where wheat blast occasionally occurs. Twenty-four isolates were analyzed, including ones from Cynodon sp. (n=1), *Digitaria* sp. (n=1), *Echinochloa* sp. (n=1), *Eleusine indica* (n=6), *Lolium multiflorum* (n=4), *Luziola* sp. (n=2), *Oryza sativa* (n=3), *Setaria italica* (n=1), *Stenotaphrum secundatum* (n=2), and Urochloa (n=3) (Table 1).

Sixteen of the 18 isolates from Uruguay either grouped according to their host-of-origin or, in the case of the *Luziola* pathogens, were representatives of a new lineage (PoLu). The only exceptions were an isolate from *Lolium multiflorum* that was recovered from a highly unusual fall-flowering plant and was placed firmly in the *P. oryzae Echinochloa* (PoEc) lineage (Fig. 1A), and an isolate from *Echinochloa* that was *P. grisea* (Fig. 1B). Critically, none of the Uruguayan isolates grouped with PoT.

To maximize our chances of gaining a representative sampling of populations found on grasses in Minas Gerais, diseased leaves (or heads in case of wheat) of grasses and wheat were collected with varying spatial and temporal separation from wheat blast-affected fields (Ascari et al., in preparation). In this manner, fungal strains were obtained from *Cenchrus echinatus, Cynodon plechtostachyus, Digitaria* spp., *Eleusine indica, Hordeum vulgare, Melinis (Rhynchelytrum) repens, Panicum maximum, Triticum aestivum* and *Urochloa* spp. (Table S1). Then, because our primary goal was to determine if the Brazilian grass-infecting *P. oryzae* are more in line with the global population than is indicated by prior research, we used sequencing of the PCR-based markers (MPG1, CH7BAC7 and CH7BAC9) to pre-screen isolates from non-wheat hosts with the goal of identifying isolates that did not appear to be members of the PoT/PoL1 lineages and were, thus, likely to represent endemic, grass-specialized lineages. Sequencing of CH7BAC7, CH7BAC9 and MPG1 PCR products for just 25 isolates revealed that 22 of them were unlikely to be PoT/PoL members, with ten of these apparently being from previously unsampled *P. oryzae* lineage and four from a different *Pyricularia* species. Genome sequences were obtained for nine isolates which variously came from *Cynodon* (n = 1), *Eleusine* (n=2), *Melinis* (n=2), *Pennisetum* (n = 2); and *Urochloa* (n = 2). In addition, we performed genome sequencing on a single isolate collected from an *Eleusine* plant growing in a heavily diseased wheat plot in an experiment station.

With just one exception, all of the isolates from Minas Gerais belonged to new lineages: the two *Eleusine* pathogens defined *P. oryzae Eleusine* lineage 3 (PoE3); CdJA159 from *Cynodon distachya* represented *P. oryzae Cynodon* lineage2 (PoC2), and MrJA49 and UbJA92 belonged to a *Melinis*-adapted lineage (PoM). The *Panicum maximum* pathogens, PmJA1 and PmJA115, showed far greater sequence divergence relative to the other isolates (~11% versus ~1%), consistent with their being members of a different species (Fig. 1B). Comparison with the NCBI database, revealed sequence identity to *Pyricularia urashimae* at a number of phylogenetic marker loci (data not shown).

The one exceptional isolate that was sampled from a wheat plot, EiJA22, grouped within the PoT lineage and was in the same haplotype group (PoT14) as (and therefore clonally related to) Ei534i, which also came from *Eleusine* growing near wheat. However, the latter isolate was collected two years earlier in Paraná state. Significantly, the only other PoT/PoL1 members that we found on grasses in S. America, were P28 and P29 from *Bromus*, and P25 from *Urochloa*, and these were all collected from grasses immediately adjacent to infected wheat.

## DISCUSSION

Current understanding on the evolution, population biology and epidemiology of wheat blast disease is largely based on the studies of Ceresini and coworkers who made several key conclusions based on the genetic relationships between *P. oryzae* found on Brazilian wheat and those found on neighboring grasses (Castroagudín et al. 2016; Ceresini et al. 2018, 2019). Throughout their work there was an implicit assumption that the isolates collected from grasses represent the populations typically found on grasses endemic to Brazil. However, several inconsistencies with prior research led us to suspect that this might not be true. Here, we report a re-analysis of their data in the context of broader community efforts which showed that all but one of the isolates used in their studies are members of the PoT or PoL1 lineages. Additionally, by surveying grass-infecting isolates collected at varying distances from wheat production areas in Brazil, we confirmed a failure to sample the true, endemic, grass-infecting populations. Furthermore, by analyzing genome sequences for select isolates, we identified a number of previously unknown *P. oryzae* lineages.

Clearly, flawed sampling was a central factor in the previous oversight, as collecting diseased grasses away from wheat fields readily turned up isolates with the expected patterns of host specialization (Ascari et al. in preparation), but another key mistake was to ignore foundational research from several groups with long histories in blast fungus research. Virtually all foregoing phylogenetic studies had shown that blast isolates from non-wheat hosts grouped according to host-of-origin (Borromeo et al. 1993; Couch and Kohn 2002; Farman 2002). So when isolates from multiple host species were found essentially to be clones of one another (e.g. Ce535i, Ei534i, Mr051i and WB037 from *Cenchrus, Eleusine, Melionis* and wheat, respectively) (Castroagudin et al. 2017; Ceresini et al. 2018, 2019) (Figs 1 and S2), this should automatically have thrown up warning flags because the sheer number of isolates that bucked the long-established trends of host specialization, by extension, would have implied that the blast populations infecting the grasses endemic to South America must behave completely differently to those found everywhere else.

Unfortunately, an immediate consequence of wheat blast having not been analyzed in the comprehensive phylogenomic framework established by the broader research community is the invalidation of most findings arising from the work. The first major conclusion to be drawn was that the wheat blast population represents a separate species (Pygt) (Castroagudín et al. 2016). When the Ceresini data were integrated into the community dataset a very different picture emerged and revealed that the seminal studies didn’t actually have the power to resolve species. This is nicely illustrated by the lone non-PoT/PoL isolate that was included in their study, Ds363 (a.k.a. 12.0.363), which was reported as having been collected from *Digitaria* spp. This appears to have been assigned to *P. grisea*, i) based on precedent (isolates from *Digitaria* historically have been *P. grisea);* and ii) because Ds363 was phylogenetically resolved from Pygt and PoO, and on an adjacent branch to the known *P. grisea* isolate, Br29. Here, we show conclusively that Ds363 belongs to *P. oryzae* and is very distantly related to Br29 (*P. grisea*) (Figure 1B). These conflicts illustrate the potential danger of trying to define species solely through empirical visualization of phylogenetic trees and emphasizes the need to incorporate statistical approaches.

Another inconsistency in the earlier studies was that Br29 consistently grouped with *P. oryzae* using phylogenomic data (Castroagudin et al. 2017; Ceresini et al. 2019) - but formed a highly diverged outgroup with the multilocus markers (Castroagudín et al. 2016). This artificial compression of phylogenetic distance for the whole genome data was largely caused by the rejection of SNPs that did not meet an arbitrary frequency threshold.

The fact that nearly all of the grass-infecting isolates included in prior studies are *bona fide* members of the PoT lineage has major implications for conclusions about wheat blast evolution, epidemiology and population biology. Previously, it was reported that “the closest relative of the wheat pathogen was found on the widely grown pasture grass *Urochloa”* (Stukenbrock and McDonald 2008) which led to the proposition that wheat blast evolved via a host jump from this grass; and that *Urochloa* probably plays a significant role in inoculum survival, production and epidemic spread. Here, it is not clear that similarity was accurately reported because the cited data (https://github.com/crolllab/wheat-blast) show that isolates from *Cenchrus* and *Eleusine* are just as closely related to the wheat blast pathogens; and the present study shows that all of these isolates, as well as others from *Elionurus* and *Melonis*, are all clones of wheat blast isolates found in their collection. So, the reason that the wheat pathogen is most closely related to isolates from *Urochloa* (and *Cenchrus, Eleusine, Elionurus* and *Melinis*), is because all of these isolates are, in fact, wheat blast pathogens. Even if we take the reported similarity at face value, the fact that the endemic *Urochloa*-infecting population was not sampled should have precluded the drawing of any conclusions regarding its possible contribution to wheat blast evolution via host jumps.

Likewise, without any information on the structure of the endemic *P. oryzae* population(s) on *Urochloa* and, especially, the relative prevalence of PoT, it seems premature to make any inferences about *Urochloa’s* role in wheat blast epidemiology. Here, it should be noted that, to date, at least 44 distinct chromosomal haplotypes have been identified within PoT (Rahnama et al. 2021; Ascari et al. in preparation), with only one of these having been found on *Urochloa* (PoT- 14). Therefore, at present, it would appear that wheat blast is more a source of inoculum for infection of *Urochloa* than vice versa.

Ceresini and coworkers found gametic equilibrium within their collection of isolates and reported this as evidence of significant gene flow between fungal isolates from the two host groups (wheat/grasses) (Ceresini et al. 2019). With the discovery that virtually all of their isolates are members of the PoT and PoL1 populations whose origins were founded through recent and abundant recombination in a multi-hybrid swarm (Rahnama et al. in review), any conclusions regarding contemporary gene flow between, or within, populations are now irrelevant. Therefore, whether or not the wheat blast population is undergoing recombination with members of grass-infecting populations remains an open question.

Lastly, Ceresini and coworkers argued that wheat blast is not a host-specialized pathogen and, therefore, “the hypothesis of grass-specific populations for the overall *P. oryzae* species complex is falsified” (Ceresini et al. 2019), so that *“P. oryzae* may not provide a suitable model for understanding the biology of *Pygt.”* Our data show that, to the contrary, wheat blast is highly host-specialized because, to date, only three out of 40 distinct PoT haplotypes have been found infecting more than one host. This is perfectly in line with what has been observed for *P. oryzae* as a whole, because while most *P. oryzae* lineages show strong host-specialization with very few exceptions, a small number of lineages show distinct non-specificity and are represented by isolates from three or more host genera - good examples are the PoEc lineage, whose members come from *Echinochloa, Leptochloa, Paspalum* and *Zea*; and PoSt comprising isolates from *Stenotaphrum, Hakonechloa, Triticum*, and *Urochloa* (Figure S2; Ascari et al. in preparation). Therefore, *P. oryzae* seems to be a perfect model for understanding wheat blast biology as it relates to host specialization.

## ACKNOWLEDGMENTS

This work was supported by the United States Department of Agriculture, Agriculture and Food Research Initiative grant 2013-68004-20378, multistate project NE1602; Agricultural Research Service project 8044-22000-046-00D; Hatch project KY012037; the National Science Foundation, MCB-1716491; and the University of Kentucky College of Agriculture Food and the Environment. Emerson M. Del Ponte was supported by the National Council for Scientific and Technological Development (CNPq) through a Productivity Research Fellowship (PQ) project 310208/2019-0, and through research grants provided by FAPEMIG. João P. Ascari was supported by CNPq through a doctoral scholarship.

---

**Table S1.**
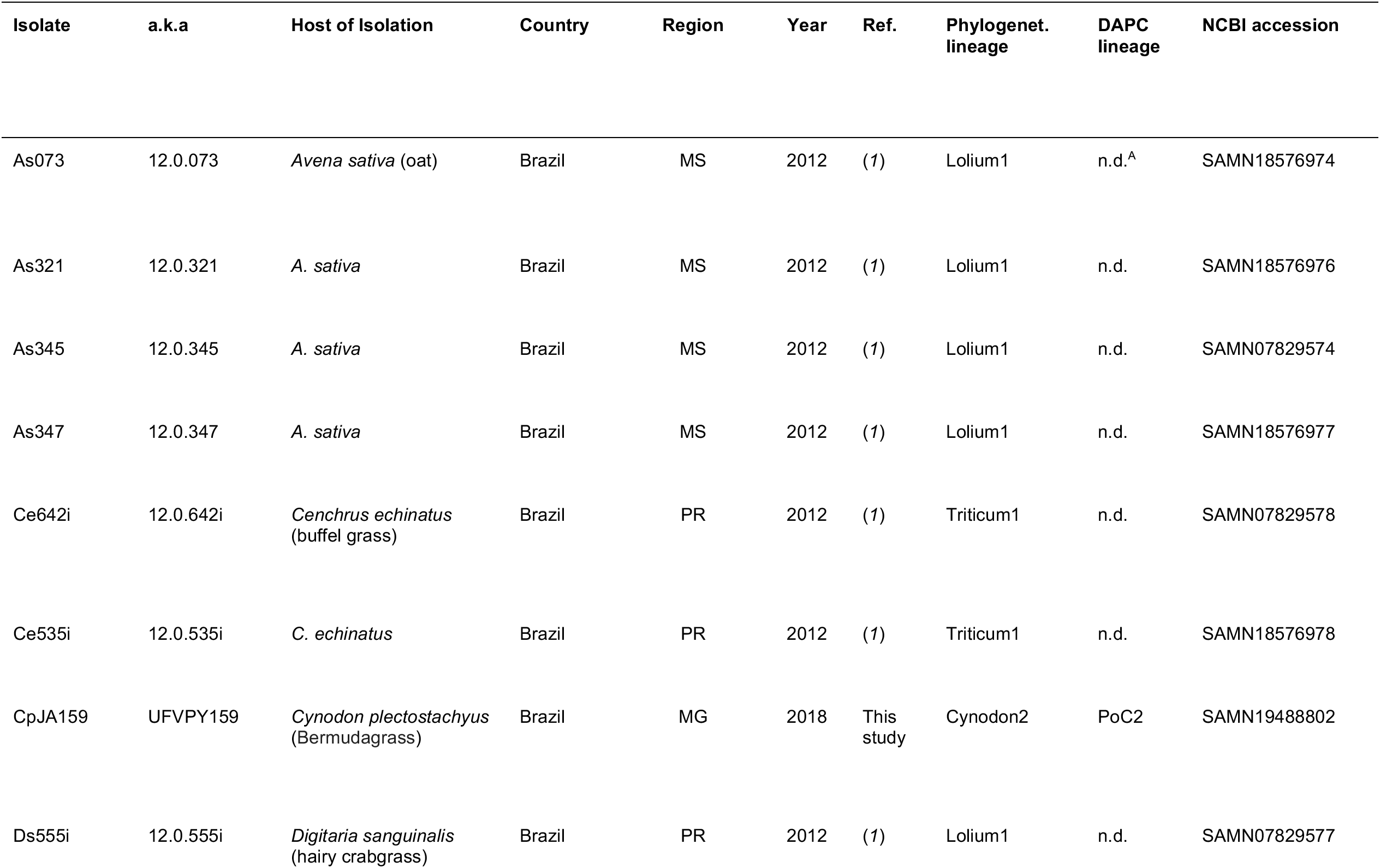
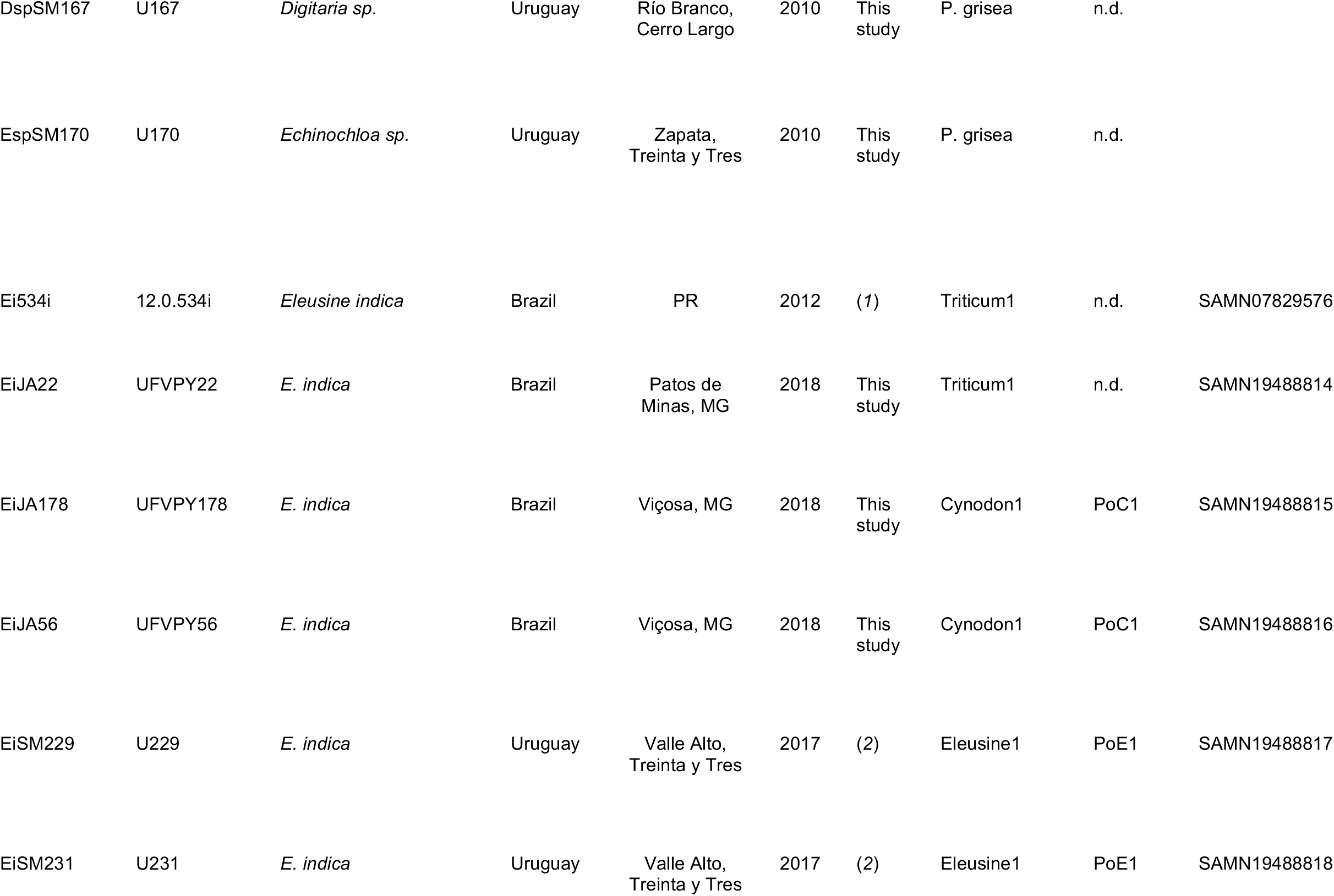
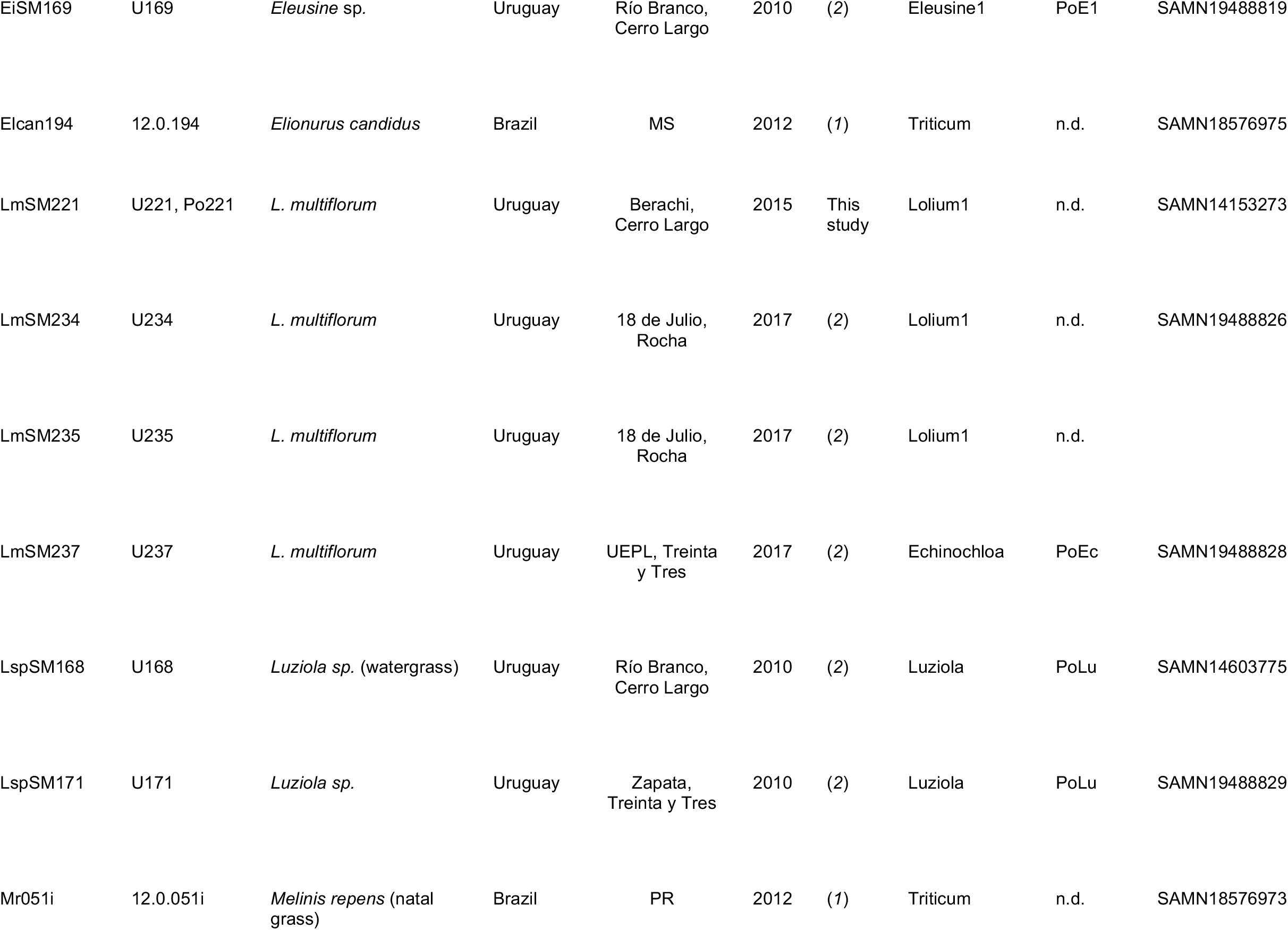
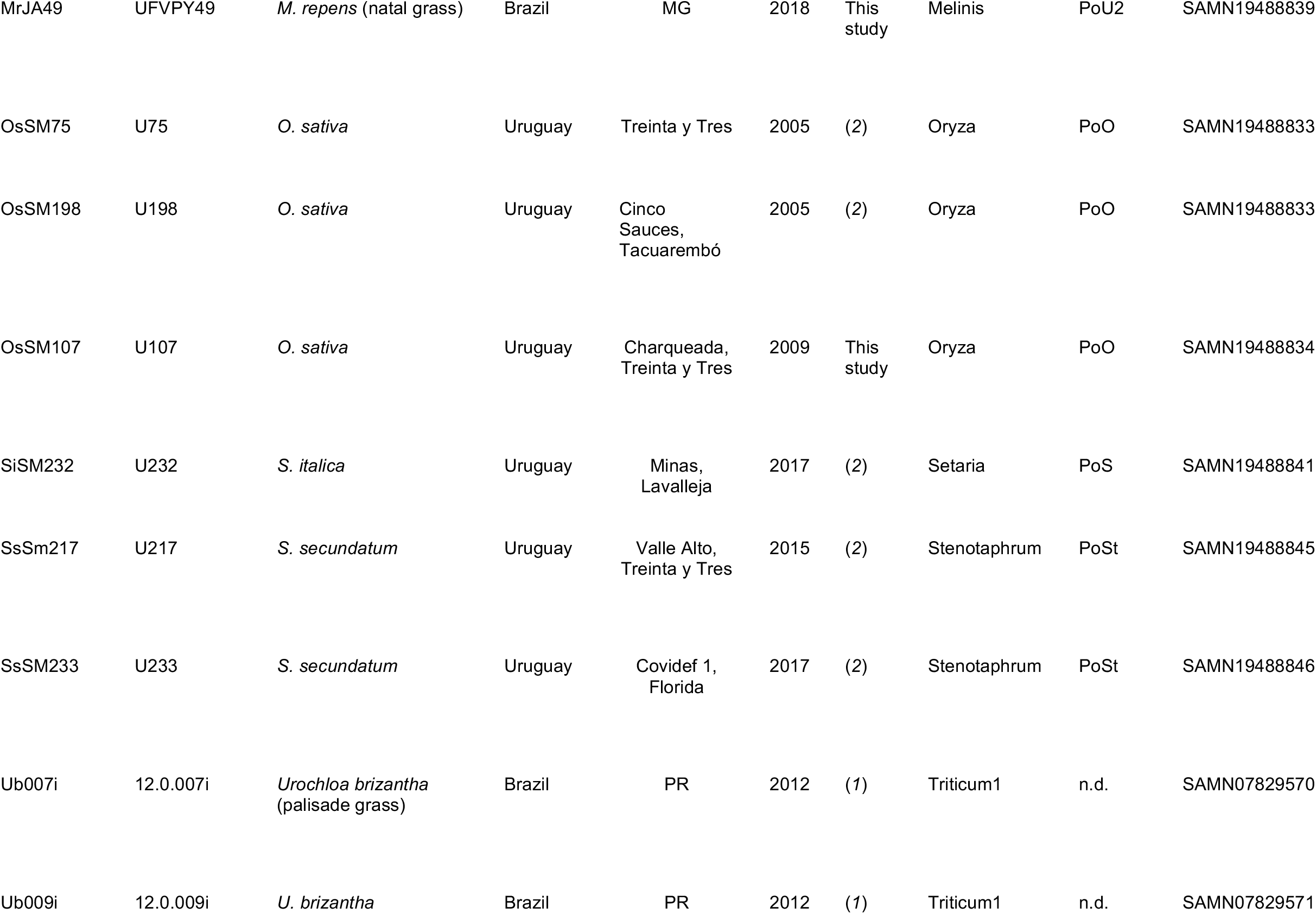
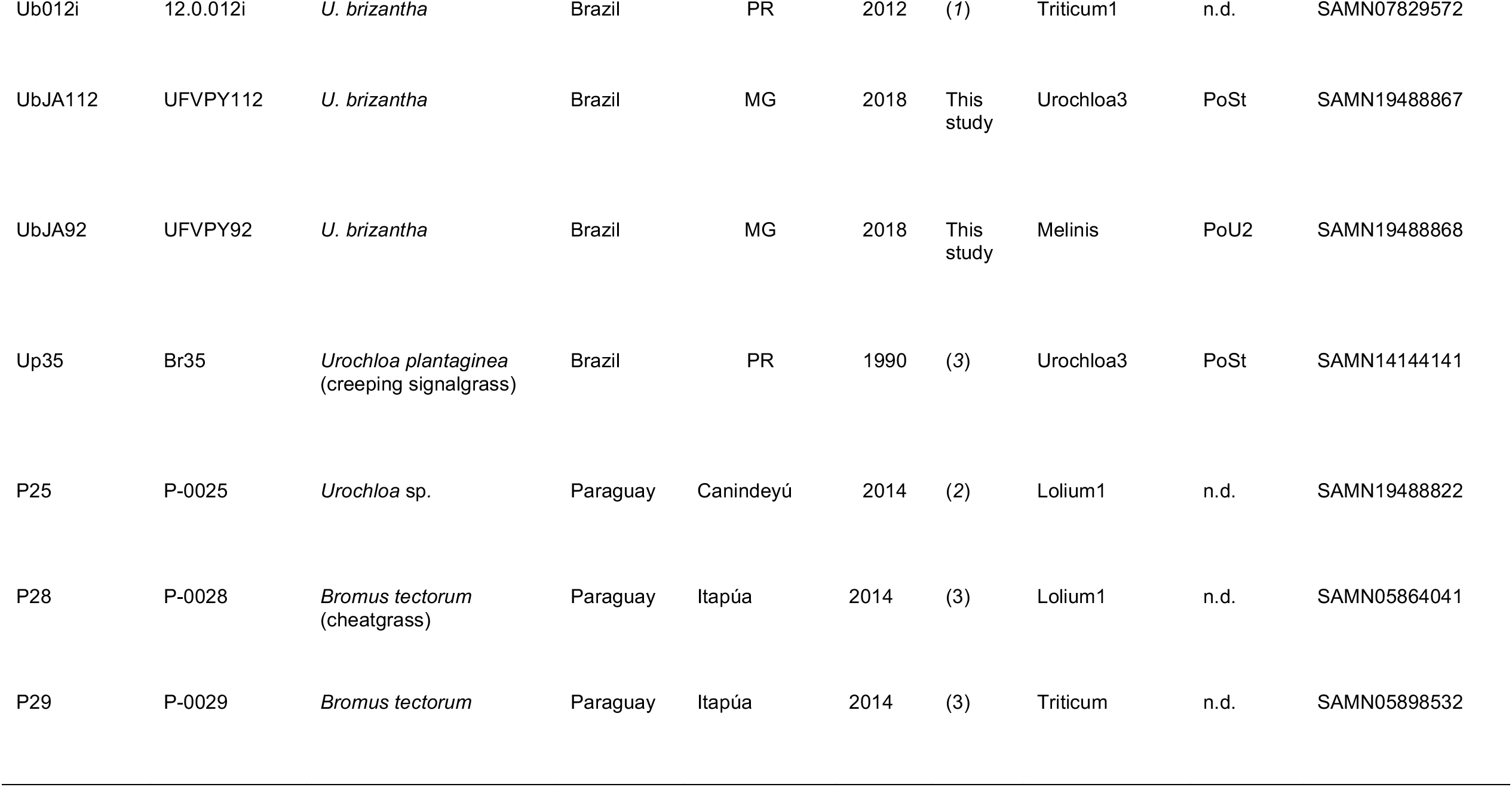
South American non-wheat blast (*non-Pyricularia oryzae* Triticum lineage) isolates (n = 41) used in this study

## Supplemental Figures

**Supplemental Figure S1.**
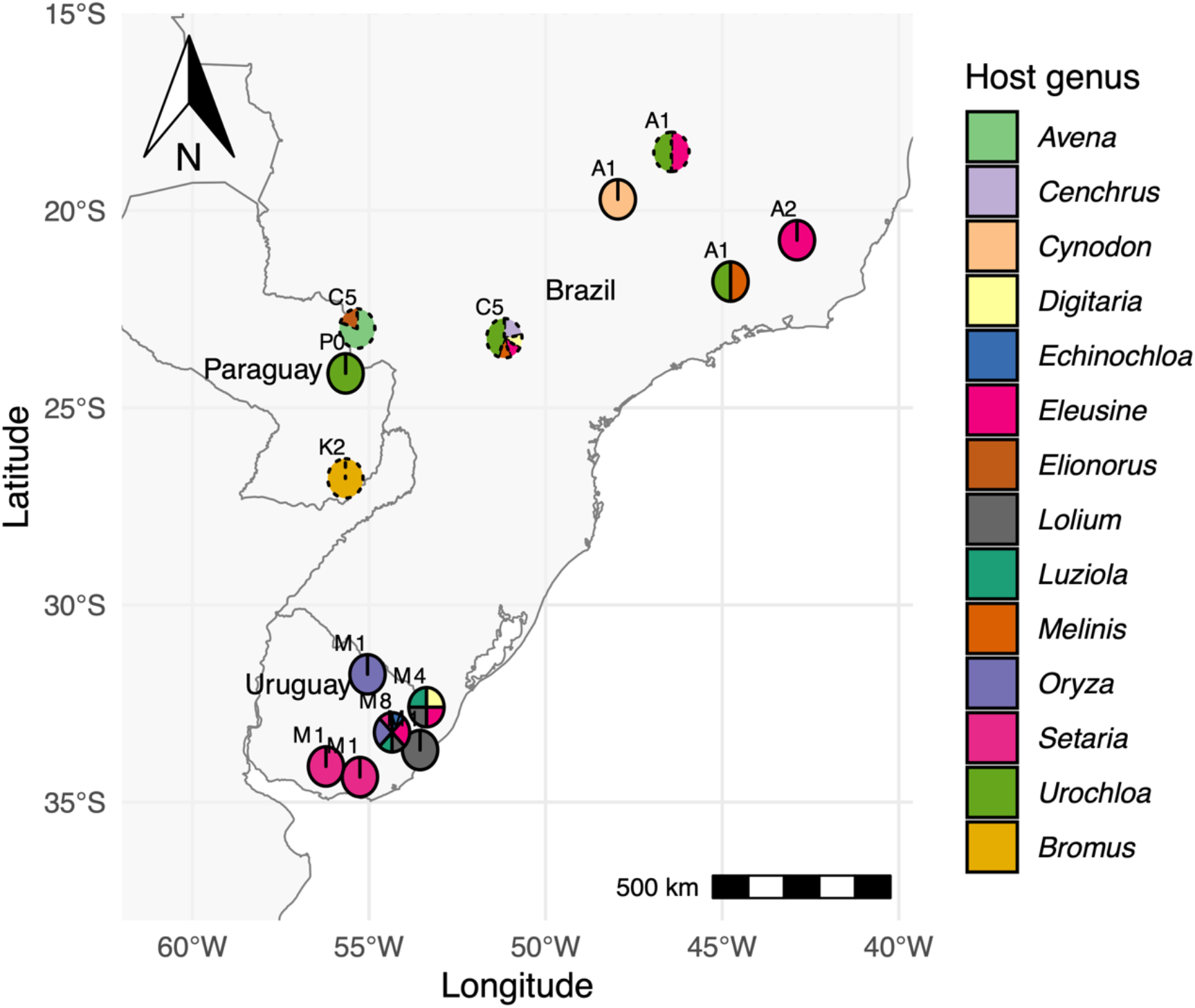
Map showing the locations where the S. American grass-infecting isolates were collected. Each sampled location is represented by a pie chart depicting the proportion of isolates from each host genus. Numbers next to each chart show the total number of isolates sampled for that location and the border style indicates the distance to the nearest wheat field (<1 km = solid line; 1-10 km = long dash; > 10 km = dotted line). Note that the isolates collected by J. Ascari and S. Martinez mostly came from regions distant from wheat fields, although the Uruguayan isolates came from regions of high *Lolium* production (and gray leaf spot incidence).

**Supplemental Figure S2.**
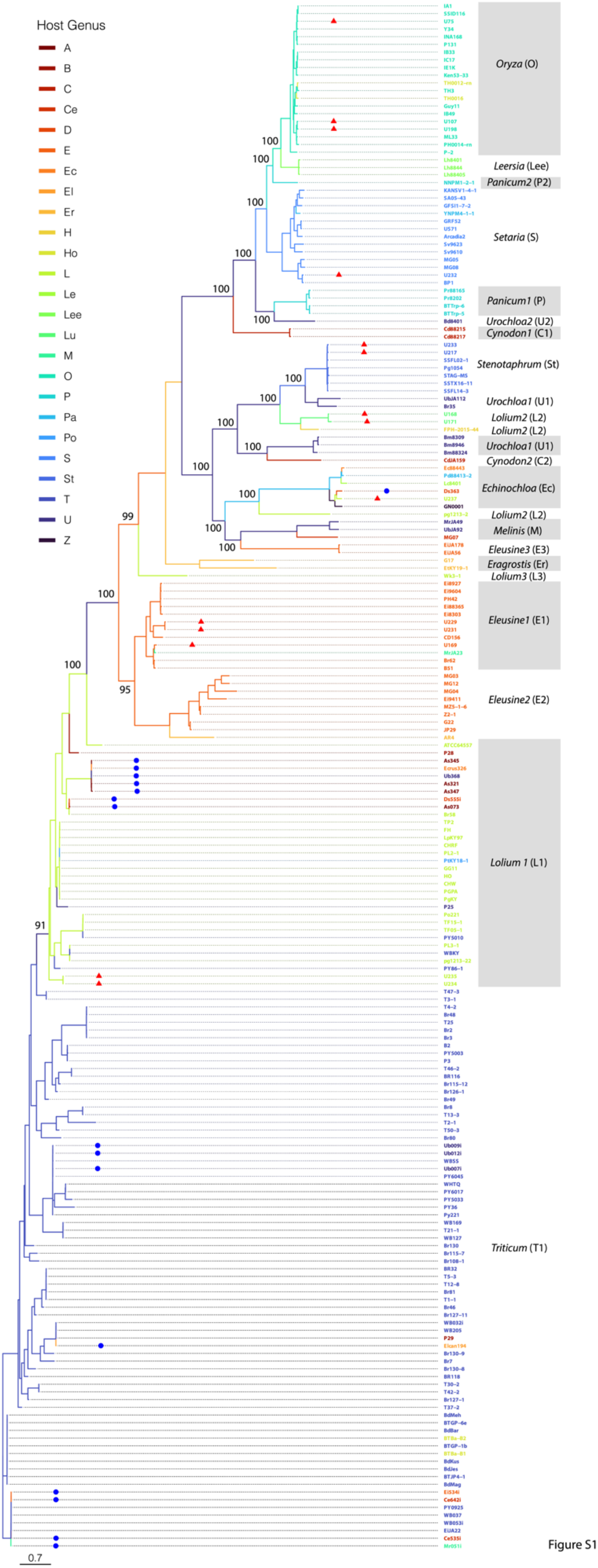
Phylogenetic relationships among isolates as revealed by Maximum Likelihood Analysis of whole genome SNP data (362,258 variant sites). Isolate names are colored according to the host of origin, while branches are colored according to phylogenetic clades as listed on the right (PoC1/2, *Cynodon1/2;* PoEc, *Echinocloa*, PoE1/2/3, *Eleusine1/2/3;* PoL1/2/3, *Lolium1/2/3;* PoLe, *Leersia;* PoM, *Melinis;* PoP, *Panicum;* PoO, *Oryza;* PoS, *Setaria;* PoSt, *Stenotaphrum;* PoT, *Triticum*, PoU1/2/3, *Urochloa1/2/3*). Host genera for which there are no distinct phylogenetic clades include *Avena* (A) and *Hordeum* (Ho) which group in PoL1 and PoO respectively. Isolates collected by the Ceresini group are highlighted with blue circles, while those from this study are marked with red triangles. Bootstrap values (100 repetitions) > 90 are presented at nodes defining the main clades.

## LITERATURE CITED

Borromeo, E. S., Nelson, R. J., Bonman, J. M., and Leung, H. 1993. Genetic differentiation among isolates of Pyricularia infecting rice and weed hosts. Phytopathology. 83:393–399.

Bolger, A. M., Lohse, M., and Usadel, B. (2014). Trimmomatic: a flexible trimmer for Illumina sequence data. Bioinformatics, 30: 2114–2120.

Bruno, A. C., and Urashima, A. S. 2001. Inter-relação sexual de Magnaporthe grisea do trigo e de outros hospedeiros. Fitopatol. Bras. 26:21–26.

Castroagudin, V., Danelli, A., Intra Moreira, S., Teodora de Assis Reges, J., Carvalho, G., Maciel, J. L. N., Bonato, A. L. V., Forcelini, C. A., Alves, E., McDonald, B. A., Croll, D., Ceresini, P. C. 2017. The wheat blast pathogen Pyricularia graminis-tritici has complex origins and a disease cycle spanning multiple grass hosts. bioRxiv. Available at: http://biorxiv.org/lookup/doi/10.1101/203455 [Accessed February 13, 2019].

Castroagudín, V. L., Moreira, S. I., Pereira, D. A. S., Moreira, S. S., Brunner, P. C., Maciel, J. L. N., Crous, P. W., McDonald, B. A., Alves, E., Ceresini, P. C. 2016. Pyricularia graminis-tritici, a new Pyricularia species causing wheat blast. Persoonia - Mol. Phylogeny Evol. Fungi. 37:199–216.

Ceresini, P. C., Castroagudín, V. L., Rodrigues, F. Á., Rios, J. A., Aucique-Pérez, C. E., Moreira, S. I., Croll, D., Alves, E., Carvalho, G., Maciel, J. L., McDonald, B. A. 2019. Wheat blast: from its origins in South America to its emergence as a global threat: Wheat Blast. Mol. Plant Pathol. 20:155–172.

Ceresini, P. C., Castroagudín, V. L., Rodrigues, F. Á., Rios, J. A., Eduardo Aucique-Pérez, C., Moreira, S. I., Alves, E., Croll, D., Maciel, J. L. 2018. Wheat blast: Past, present, and future. Annu. Rev. Phytopathol. 56:427–456.

Couch, B. C., and Kohn, L. M. 2002. A multilocus gene genealogy concordant with host preference indicates segregation of a new species, Magnaporthe oryzae, from M. grisea. Mycologia. 94:683–693.

Cruz, C. D., and Valent, B. 2017. Wheat blast disease: danger on the move. Trop. Plant Pathol. 42:210–222.

Farman, M. L. 2002. Pyricularia grisea isolates causing gray leaf spot on perennial ryegrass (Lolium perenne) in the United States: Relationship to P. grisea isolates from other host plants. Phytopathology. 92:245–254.

Galbieri, R., and Urashima, A. S. 2008. Caracterização, compatibilidade e ocorrência de reprodução sexual entre isolados de Pyricularia grisea de diferentes hospedeiros. Summa Phytopathol. 34:22–28.

Gladieux, P., Condon, B., Ravel, S., Soanes, D., Maciel, J. L. N., Nhani, A., Chen, L., Terauchi, R., Lebrun, M. H., Tharreau, D., Mitchell, T., Pedley, K. F., Valent B., Talbot, N. J., Farman, M., Fournier, E. 2018. Gene flow between divergent cereal- and grass-specific lineages of the rice blast fungus Magnaporthe oryzae mBio. 9(1). DOI: 10.1128/mBio.01219-17

Igarashi, S., Utiamada, C. M., Igarashi, L. C., Kazuma, A. H., Lopes, R. S. 1986. Occurrence of Pyricularia sp. in wheat (Triticum aestivum L.) in the State of Parana, Brazil. Fitopatol. Bras. 11:351–352.

Kumar, S., Stecher, G., Li, M., Knyaz, C. and Tamura, K., 2018. MEGA X: molecular evolutionary genetics analysis across computing platforms. Mol. Biol. Evo. 35(6), p.1547.

Maciel, J. L. N., Ceresini, P. C., Castroagudin, V. L., Zala, M., Kema, G. H. J., and McDonald, B. A. 2014. Population structure and pathotype diversity of the wheat blast pathogen Magnaporthe oryzae 25 years after its emergence in Brazil. Phytopathology. 104:95–107.

Malaker, P. K., Barma, N. C. D., Tiwari, T. P., Collis, W. J., Duveiller, E., Singh, P. K., Joshi, A. K., Singh, R. P., Braun, H. J., Peterson, G. L., Pedley, K. F., Farman, M. L., Valent, B. 2016. First Report of wheat blast caused by Magnaporthe oryzae pathotype Triticum in Bangladesh. Plant Dis. 100:2330.

Milgroom, M. G. 2017. Population Biology of Plant Pathogens: Genetics, Ecology, and Evolution. The American Phytopathological Society.

Rahnama, M., Condon, B., Ascari, J. P., Dupuis, J. R., Del Ponte, E. M, Pedley, K. F., Martinez, S., Valent, B., Farman, M. L. 2021. Recombination of standing variation in a multi-hybrid swarm drove adaptive radiation in a fungal pathogen and gave rise to two pandemic plant diseases. BiorXiv:2021.11.24. DOI: 10.1101/2021.11.24.469688v1

Singh, P. K., Gahtyari, N. C., Roy, C., Roy, K. K., He, X., Tembo, B., Xu., K., Juliana, P., Sonder, K., Kabir, M. R., Chawade, A. 2021. Wheat blast: A disease spreading by intercontinental jumps and its management strategies. Front. Plant Sci. 12:1–21.

South, A. 2021. rnaturalearth: World Map Data from Natural Earth. Available at: https://github.com/ropensci/rnaturalearth.

South, A. 2022. rnaturalearthhires: High Resolution World Vector Map Data from Natural Earth used in rnaturalearth. Available at: https://github.com/ropensci/rnaturalearthhires.

Stukenbrock, E. H., and McDonald, B. A. 2008. The origins of plant pathogens in agroecosystems. Annu. Rev. Phytopathol. 46:75–100.

Tembo, B., Mulenga, R. M., Sichilima, S., M’siska, K. K., Mwale, M., Chikoti, P. C., Singh, P. K., he, X., Pedley, K. F., Peterson, G. L., Singh, R. P., Braun, H. J. 2020. Detection and characterization of fungus (Magnaporthe oryzae pathotype Triticum) causing wheat blast disease on rain-fed grown wheat (Triticum aestivum L.) in Zambia. PLOS ONE. 15:1–10.

Urashima, A. S., Igarashi, S., and Kato, H. 1993. Host range, mating type, and fertility of Pyricularia grisea from wheat in Brazil. Plant Dis. 77:1211–1216.

Valent, B., Crawford, M. S., Weaver, C. G., and Chumley, F. G. 1984. Genetic studies of fertility and pathogenicity in Magnaporthe grisea (Pyricularia oryzae). United States: N. p.

Valent, B., Farman, M., Tosa, Y., Begerow, D., Fournier, E., Gladieux, P., Islam, M. F., Kamoun, S., Kemler, M. Kohn, L. M., Lebrun, M. H., Stajich, J. E., Talbot, N. J., Terauchi, R. Tharreau, D., Zhang, N. 2019. Pyricularia graminis-tritici is not the correct species name for the wheat blast fungus: response to Ceresini et al. (MPP 20:2). Mol. Plant Pathol. 20:173–179.

Yu, G. 2021. scatterpie: Scatter Pie Plot. R package version 0.1.7. Available at: https://CRAN.R-project.org/package=scatterpie.

Yu, G., Smith, D. K., Zhu, H., Guan, Y., and Lam, T. T.-Y. 2017. ggtree: an r package for visualization and annotation of phylogenetic trees with their covariates and other associated data. Methods Ecol. Evol. 8:28–36.

